# The foliar endophyte *Phialocephala scopiformis* DAOMC 229536 secretes enzymes supporting growth on wood as sole carbon source

**DOI:** 10.1101/354365

**Authors:** Jennifer M. Bhatnagar, Grzegorz Sabat, Daniel Cullen

## Abstract

The conifer needle endophyte, *Phialocephala scopiformis*, was cultivated in media containing ground *Pinus contorta* wood as sole carbon source. After five and seven days growth, concentrated extracellular fluids were subjected to LC-MS/MS analyses. A total of 590 proteins were identified of which 99 were assigned to glycoside hydrolase families within the Carbohydrate Active Enzyme (CAzyme) system. Multiple isozymes of exo-and endo-acting cellulases were among the most abundant proteins, and oxidative degradation of cellulose was supported by the presence of lytic polysaccharide monooxygenases, glucooligosaccharide oxidase and cellobiose dehydrogenase. Oxidoreductases were also plentiful and included GMC oxidoreductases, alcohol dehydrogenases, laccases, copper radical oxidases, tyrosinases and catalase. The expression and diversity of extracellular oxidoreductases indicates a capacity to metabolize alcohols and aromatic compounds.

## Introduction

Microbial degradation of wood, a pivotal process in terrestrial carbon cycling, involves the activities of a diverse array of fungi and bacteria (1–3). A common assumption is that wood is decomposed by free-living saprotrophic fungi, primarily wood rot species in the fungal phylum Basidiomycota (4). Among these wood rot fungi, certain members of the subdivision Agaricomycotina, particularly the white-rot fungi, efficiently depolymerize and mineralize lignin. Extracellular class II peroxidases, and perhaps laccases, have been repeatedly implicated in ligninolyis by these Basidiomycetes. Low efficiency lignin degraders *Jaapia argillacea* and *Botryobasidium botryosum* lack class II peroxidases (5) and are unable to oxidize lignin model compounds. Still, these slow-growing fungi erode wood cell walls and lignin is removed. Other members of the Agaricomycotina, collectively referred to as brown rot fungi, rapidly depolymerize the cellulose within plant cell walls while leaving the lignin as a modified residue (6).

While most research has focused on the lignin-degrading abilities of the Basidiomycota, evidence is emerging that fungal species in the other dominant fungal phylum, Ascomycota, may also play a significant role in decay of lignified wood. Evidence supports lignin degradation by several members of the Ascomycete family, Xylariaceae. This capability was most convincingly demonstrated for *Daldinia concentrica* which efficiently degrades hardwood lignin as well as 14C-labeled synthetic lignins (i.e. DHPs) (7, 8). Other Ascomycete wood decayers, collectively referred to as ‘soft rot’, have received relatively little attention (9). Some capacity to degrade cellulose and hemicellulose is presumed, but their role in ligninolysis, if any, is unknown. This is a significant knowledge gap, as the Ascomycetes are a diverse and abundant fungal group on land.

In particular, Ascomycete fungi comprise most fungal species that associate with live plant tissues, including endophytes (10). Dark septate endophytes (DSEs) are a large and widely distributed group considered nutritionally distinct from the free-living wood decay saprophytes (11). However, accumulating evidence shows that many DSEs exhibit unexpectedly broad substrate utilization patterns (12–16) and gene complements (17, 18) suggesting significant contributions to carbon cycling via latent saprotrophy. Nevertheless, the enzymes these organisms produce, and biochemical mechanism(s) they use to depolymerize plant biopolymers, have not been identified.

*Phialocephala scopiformis* and *Phialocephala subalpina* are well known endophytes of conifer needles and roots, respectively. Nursery inoculation of conifer seedlings with *P. scopiformis* limits spruce budworm damage, and persists in mature trees (19). Recently sequenced, the genome sizes/predicted gene numbers for *P. scopiformis* and closely related *P. subalpina*, are estimated to be 48.9Mb/18,613 (20) and 69.7 Mb/20,173 (17), respectively. Schlegel et al (17) noted an impressive repertoire of predicted genes in *P. subalpina*, particularly the diversity and number of those encoding Carbohydrate Active Enzymes (CAzymes; (21)). The authors suggested that this gene expansion might confer the capability of saprophytic growth utilizing plant-derived substrates. Metatranscriptome examination of extensively decayed Pinus contorta logs (SRA accessions SAMN07573382, SAMN07573383 and SAMN07573384) have shown that transcripts most closely related to *P. scopiformis*, followed by *P. subalpina*, constitute the most abundant Ascomycetes present.

Here, we show that the previously sequenced endophyte of conifer needles, *P. scopiformis* DAOMC 229536, is indeed capable of utilizing *P. contorta* wood as a sole carbon source and, in doing so, secretes an array of hydrolytic and oxidative enzymes.

## Materials and methods

*P. scopiformis* DAOMC 229536 was obtained from the Canadian Collection of Fungal Cultures. Two-liter flasks containing 250 ml of basal salt media (22) were supplemented with 1.25 g of *P. contorta* wood ground in a Wiley-mill (30 mesh screen) as the sole carbon source. Cultures were inoculated with mycelium scraped from the surface of malt extract agar (2% w/w malt extract, 2% glucose w/w, 0.5% peptone, 1.5% agar) and placed on a rotary shaker (150 RPM). After five and seven days incubation at 22-24C, cultures were filtered successively through Miracloth (Calbiochem) and Whatman filter paper #50 and #541. Trichloroacetic acid was added 10% (wt/vol) to the cold filtrate which was stored overnight at 0C. The precipitate was centrifuged at 12,000 × g for 20 min and the pellet washed three times in cold acetone before air drying. Total proteins from the pellet fraction were further purified via methanol/chloroform/water partitioning, where chloroform and methanol were added to pellet first, followed by water and allowed to partition with protein interphase formed between polar and non-polar fraction. After multiple methanol washes, these finely purified protein preps where ultimately resolubilized in 8M Urea / 50mM NH_4_HCO_3_ (pH8.5) / 1mM TrisHCl and a fraction taken for protein concentration determination (PIERCE™ 660nm Protein Assay kit, ThermoFisher Scientific). NanoLC-MS/MS was used to identify proteins in culture filtrates as described (23–25). In short, equal amounts of total protein per sample were trypsin/LysC digested, OMIX C18 SPE purified (Agilent Technologies) and finally 2μg loaded for nanoLC-MS/MS analysis using an Agilent 1100 nanoflow system (Agilent Technologies) connected to a hybrid linear ion trap-orbitrap mass spectrometer (LTQ-Orbitrap Elite™, ThermoFisher Scientific) equipped with an EASY-Spray™ electrospray source. Chromatography of peptides prior to mass spectral analysis was accomplished using capillary emitter column (PepMap^®^ C18, 3μM, 100Å, 150×0.075mm, ThermoFisher Scientific) onto which 2μl of purified peptides was automatically loaded. NanoHPLC system delivered solvents A: 0.1% (v/v) formic acid, and B: 99.9% (v/v) acetonitrile, 0.1% (v/v) formic acid at 0.50 μL/min to load the peptides (over a 30 minute period) and 0.3μl/min to elute peptides directly into the nano-electrospray with gradual gradient from 3% (v/v) B to 20% (v/v) B over 154 minutes and concluded with 12 minute fast gradient from 20% (v/v) B to 50% (v/v) B at which time a 5 minute flash-out from 50-95% (v/v) B took place. As peptides eluted from the HPLC-column/electrospray source survey MS scans were acquired in the Orbitrap with a resolution of 120,000 followed by MS2 fragmentation of 20 most intense peptides detected in the MS1 scan from 380 to 1800 m/z; redundancy was limited by dynamic exclusion. Raw MS/MS data were converted to mgf file format using MSConvert (ProteoWizard: Open Source Software for Rapid Proteomics Tools Development) for downstream analysis. Resulting mgf files were used to search against Phialocephala scopiformis protein database containing 18,573 entries via the JGI portal (https://genome.jgi.doe.gov/portal/pages/dynamicOrganismDownload.jsf?organism=Phisc1) with a list of common lab contaminants and decoy sequences to establish False Discovery Rate (37,222 total entries) using in-house *Mascot* search engine 2.2.07 [Matrix Science] with variable methionine oxidation, asparagine and glutamine deamidation plus fixed cysteine carbamidomethylation. Scaffold (version Scaffold_4.7.5, Proteome Software Inc., Portland, OR) was used for spectral based quantification and to validate MS/MS peptide and protein identifications. Peptide identifications were accepted if they could be established at greater than 80.0% probability to achieve an FDR less than 1.0% by the Scaffold Local FDR algorithm. Protein identifications were accepted if they could be established at greater than 99.0% probability to achieve an FDR less than 1.0% and contained at least 2 identified peptides. Protein probabilities were assigned by the Protein Prophet algorithm (26). Proteins that contained similar peptides and could not be differentiated based on MS/MS analysis alone were grouped to satisfy the principles of parsimony. The raw mass spectrometry data was deposited to Chorus repository (chorusproject.org, ID#1478) for public sharing and visualization.

Proteins were classified by blastp queries of NCBI NR and EMBL-EBI predictions of Interpro domains and secretion signals. Carbohydrate active enzyme (CAzyme) family assignments were as described (21)

## Results and discussion

We found that the spruce endophyte *P. scopiformis* DAOMC 229536 will utilize lodgepole pine wood as a sole carbon source and, to do so, a diverse set of extracellular enzymes are expressed. Peptides corresponding to 590 proteins were identified from this species and broadly classified (Fig 1). Those categorized as CAzymes included glycoside hydrolases (GHs), auxiliary activities (AAs) and carbohydrate esterases (CEs), and accounted for 24% of the total proteins. Highly expressed ribosomal and ‘housekeeping’ proteins involved in central metabolism made up 56% of the total. These are typically intracellular proteins, and their detection is likely a result of hyphal lysis. Although function could not be predicted, the 84 sequences classified as ‘hypothetical’ were generally conserved in other fungi and 32 featured clear secretion signals. One of these, model Phisc_220767, had the highest protein abundance score in both samples. Interestingly, the sequence contains InterPro domain IPR021054 that is predicted to bind hydrophobic surfaces. In *Aspergillus oryzae* cultures, binding to the polymer polybutylene succinate-co-adipate mobilizes degradation by cutinase (27). In this connection, we also detected a cutinase protein in the *P. scopiformis* cultures (Phisc_572142). Most (82%) of the identified proteins were present at both time points, and emPAI values were generally highest after seven days growth. Detailed information for all 590 proteins is listed in supporting information (S1 File).

**Fig. 1.**
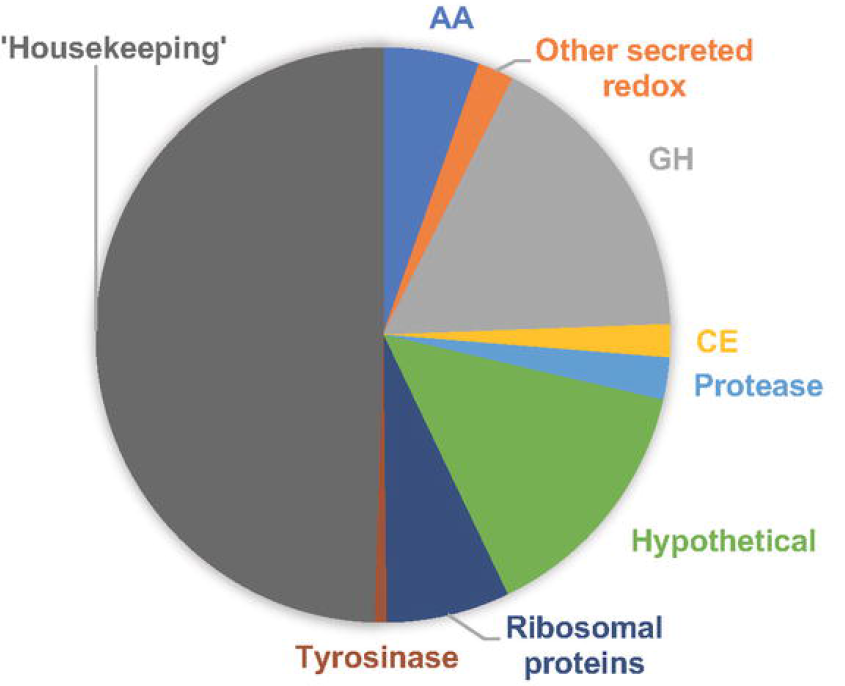
Distribution of 590 *P. scopiformis* proteins detected in media containing ground lodgepole pine as the sole carbon source after five or seven days incubation. Of these, 482 proteins were identified at both time points, 10 were exclusively detected at day five and 98 were found only in day seven cultures (see S1 File for details).

The *P. scopiformis* genome is predicted to encode 387 glycoside hydrolases (https://genome.jgi.doe.gov/mycocosm/annotations/browser/cazy/summary;f4VTMj?p=Phisc1) and 99 were identified here. These were distributed among 39 families or subfamilies (Fig 2). For most families, a fraction of the predicted proteins was observed, but all representatives of GH36, GH39, GH5_7, GH53, GH54 and GH74 were identified. No clear relationship was observed between the number of genes per family and their detection (Fig 2) or their abundance (Table 1). Cellulases were among the most abundant enzymes and included the ‘exo’-acting cellobiohydrolases (GH6s, GH7s), endoglucanses (GH5_5) and beta-glucosidase (GH3) (Table 1). All had secretion signals and family one carbohydrate binding modules (CBM1s) were predicted in six of the cellulases. Other GHs were more likely involved in the degradation of hemicellulose which, in conifers such as lodgepole pine, is primarily composed of O-acetylgalactoglucomannans and nonacetlylated arabinoglucuronoxylans (28). The identified hydrolases (Table 1) are well suited for debranching and depolymerizing these complex substrates (reviewed in (29)). Inspection of the 14 predicted GH78s, including the three identified by mass spectrometry, failed to identify a feruloyl esterase domain in addition to the expected rhamnosidase. Such a bifunctional enzyme has been implicated in lignocellulose degradation by the soft rotter *Xylaria polymorpha* (30). No secretion signal was predicted for an Alpha-arabinofuranosidase (Phisc_760840) and Alpha-galactosidase (Phisc_659201), but alternative models Phisc_597046 and Phisc_659199, respectfully, produced high-confidence eukaryote signals.

**Fig. 2.**
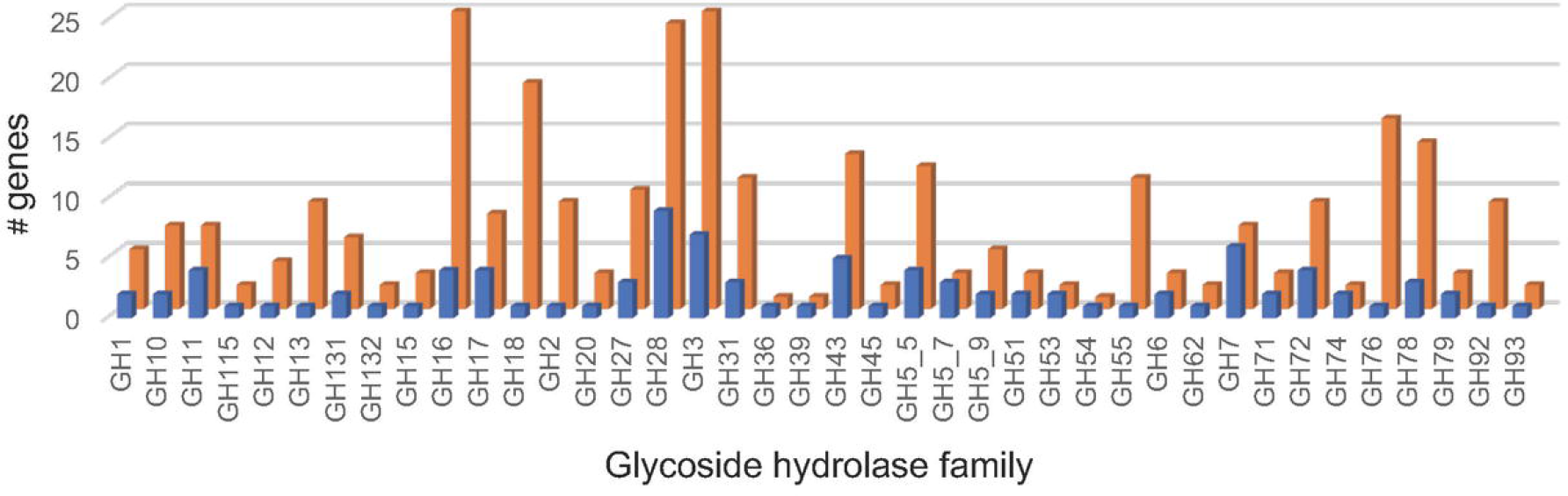
Predicted genes and LC-MS/MS detected glycoside hydrolases. The number of predicted genes https://genome.jgi.doe.gov/Phisc1/Phisc1.home.html and detected proteins are shown by brown and blue bars, respectively.

**Table 1.** Proteins classified as CAzyme Glycoside Hydrolases and Carbohydrate Binding Modules^1^.

| Protein | 5 Day | 7 Day | CAzyme | Putative function | comment |
| --- | --- | --- | --- | --- | --- |
| 668883 | 14.220 | 27.784 | GH7-CBM | Cellobiohydrolase (CBH1) | SS |
| 709446 | 9.673 | 22.500 | GH5_5-CBM1 | Endo-1,4-beta-glucanase | SS |
| 573362 | 5.670 | 21.688 | GH54 | Alpha-N-arabinofuranosidase | SS |
| 650591 | 4.158 | 13.740 | GH7 | Cellobiohydrolase (CBH1) | SS |
| 703845 | 2.129 | 6.525 | GH5_5 | Endo-1,4-beta-glucanase | SS |
| 749482 | 0.791 | 4.340 | GH5_7 | Endo-beta-1,4-mannanase | SS |
| 617769 | 1.831 | 4.083 | GH27 | Alpha galactosidase | SS |
| 760840 | 1.464 | 3.633 | GH62 | Alpha-L-arabinofuranosidase |  |
| 591570 | 1.418 | 3.425 | GH15-CBM20 | 1,4-alpha-D-glucan glucohydrolase | SS |
| 785300 | 1.254 | 3.213 | GH39 | Beta-xylosidase | SS |
| 580672 | 0.715 | 2.853 | GH5_7-CBM1 | Endo-beta-1,4-mannanase | SS |
| 118029 | 0.976 | 2.356 | GH93 | Alpha-L-arabinofuranosidase | SS |
| 613046 | 1.917 | 2.149 | GH6 | Cellobiohydrolase (CBH2) | SS |
| 706718 | 0.717 | 1.950 | GH6-CBM1 | Cellobiohydrolase (CBH2) | SS |
| 706461 | 0.484 | 1.864 | GH7-CBM | Cellobiohydrolase (CBH1) | SS |
| 574388 | 0.241 | 1.818 | GH78 | Alpha-L-rhamnosidase | SS |
| 639775 | 0.348 | 1.641 | GH53 | Arabinogalactan endo-1,4-beta-galactosidase | SS |
| 539647 | 0.316 | 1.616 | GH11 | Endo-1,4-beta-xylanase | SS |
| 701593 | 0.536 | 1.475 | GH13-CBM20 | Alpha-amylase | SS |
| 783091 | 1.193 | 1.372 | GH28 | Endopolygalacturonase | SS |
| 595918 | 0.928 | 1.347 | GH7 | Cellobiohydrolase (CBH1) | SS |
| 593362 | 0.110 | 1.312 | GH131 | Exo/Endo glucanase | SS |
| 725142 | 0.447 | 1.296 | GH51 | Alpha-N-arabinofuranosidase | SS |
| 659201 | 0.791 | 1.261 | GH27 | Alpha-galactosidase |  |
| 679634 | 0.144 | 1.096 | GH5_9 | Glucan 1,3-beta-glucanase | SS |
| 687038 | 0.039 | 1.080 | GH3 | Beta-glucosidase | SS |
| 756372 | 0.130 | 1.078 | GH11 | Endo-1,4-beta-xylanase | SS |
| 588206 | 0.242 | 1.061 | GH10-CBM1 | Endo-1,4-beta-xylanase | SS |
| 758257 | 0.719 | 0.969 | GH11 | Endo-1,4-beta-xylanase | SS |
| 709032 | 0.287 | 0.961 | GH53 | Endo-1,4-beta-galactanase | SS |

<sup>1</sup>Thirty CAzymes (31) with highest exponentially modified protein abundance indexes (emPAI) (32) after 5 and 7 day growth on ground lodgepole pine. Abbreviations: GH, glycoside hydrolase; CBM, carbohydrate binding module; SS, secretion signal

In addition to conventional hydrolases long known to degrade cellulose and hemicellulose, an array of extracellular oxidative enzymes and hypothetical proteins were unambiguously detected. The precise role of some of these enzymes remains uncertain, but the impressive diversity suggests complex strategies involved in lignocellulose degradation (Table 2). A glucose-methanol-choline (GMC) oxidoreductase (Phisc_171447) was the second most abundant protein in cultures. A putative alcohol oxidase, the sequence is highly conserved among a wide range of taxa. Other AA3 family members include putative glucose oxidases which, together with the copper radical oxidases (AA5_1s), may generate extracellular H_2_O_2_. Peroxide production is also observed in brown rot Basidiomycetes in which cellulose depolymerization is generally attributed to hydroxyl radical production via Fenton chemistry (33–39). Benzoquinone reductase (Phisc_680385) and oxalate decarboxylase (Phisc_490473; Table 3) have also been implicated in this wood decay process (38, 40). In addition to the glycoside hydrolases (Table 1) and the possibility of Fenton-generated hydroxyl radical, *P. scopiformis* may directly attack cellulose (41, 42) and xylans (43, 44) with Lytic polysaccharide monoxygenases (LPMO; AA9s), Cellobiose dehdogenase (CDH; AA3_1) and Glucooligosaccharide oxidases (GOOX; AA7). Peptides corresponding to nine model LPMO proteins were detected and, in line with the abundance of AA3s and AA5s, these copper enzymes are now known to be dependent upon H_2_O_2_ (45, 46). In short, *P. scopiformis* employs both hydrolytic and oxidative strategies for the utilization of *P. contorta* wood.

**Table 2.**
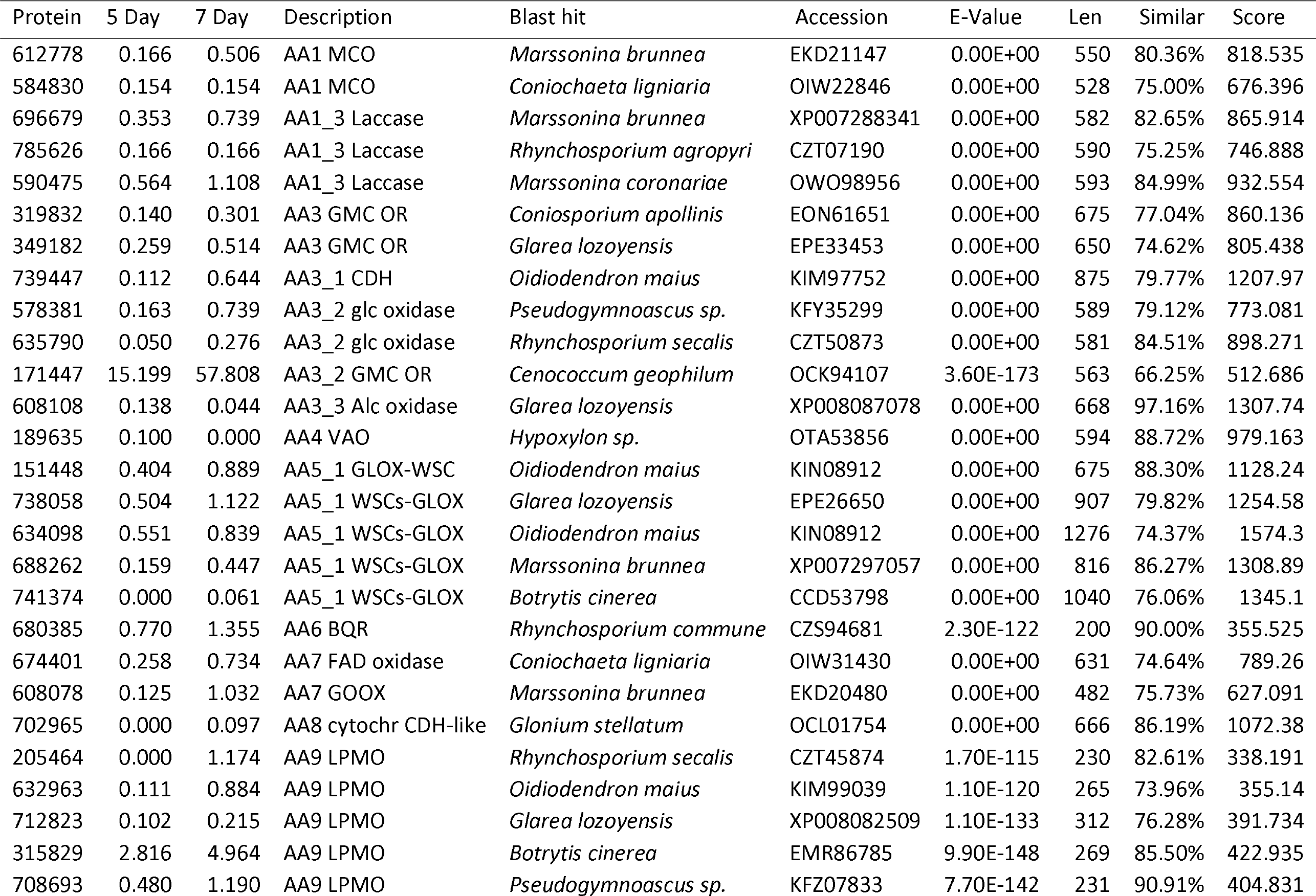
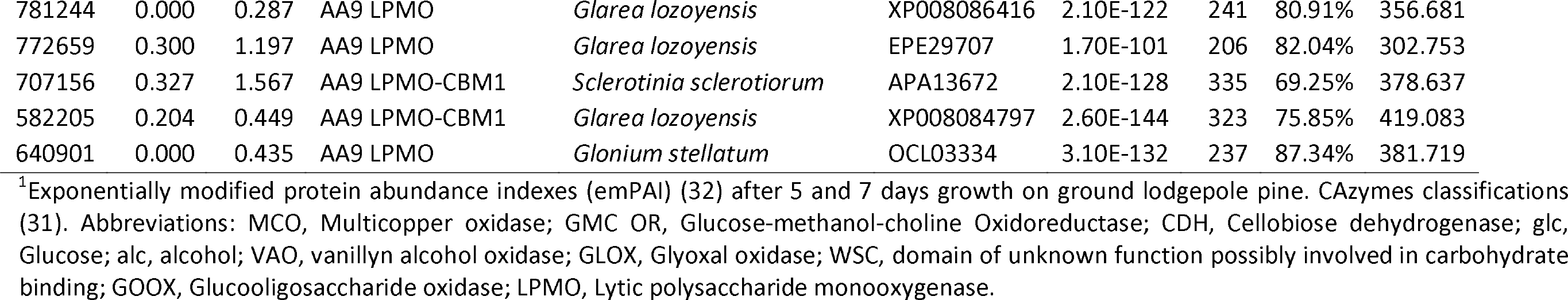
Proteins classified as members of Auxiliary Activities families^1^.

**Table 3.** Mass spectrometry identified proteins with predicted secretion signals and redox activitiy^1^.

| Protein | 5 Day | 7 Day | Putative function | Blast Hit | Accession | E-Value | Score |
| --- | --- | --- | --- | --- | --- | --- | --- |
| 729673 | 0.304 | 0.489 | Heme peroxidase-WSCs | *Glarea lozoyensis* | XP_008077688 | 0.00E+00 | 682.945 |
| 766309 | 0.278 | 0.445 | Catalase | *Botrytis cinerea* | AAK77951 | 0.00E+00 | 1135.55 |
| 706407 | 0.000 | 0.146 | FAD-binding oxidoreductase | *Botrytis cinerea* | EMR83325 | 0.00E+00 | 884.789 |
| 490473 | 1.010 | 1.761 | Oxalate decarboxylase | *Marssonina coronariae* | OWO98734 | 0.00E+00 | 749.969 |
| 576987 | 0 | 0.148 | Chloroperoxidase | *Marssonina coronariae* | OWP06232 | 0.00E+00 | 687.567 |
| 588357 | 0.145 | 0.226 | Chloroperoxidase | *Rhynchosporium secalis* | CZT46035 | 0.00E+00 | 737.643 |
| 54274 | 0.588 | 1.614 | Quinoprotein-ADH-like WSC | *Marssonina brunnea* | EKD19016 | 4.30E-179 | 569.311 |
| 682730 | 0.799 | 1.275 | Quinoprotein-ADH-like WSC | *Macrophomina phaseolina* | EKG22475 | 0.00E+00 | 1586.24 |
| 447552 | 0.065 | 0.159 | Quinoprotein-ADH-like WSC | *Sclerotinia sclerotiorum* | APA14815 | 0.00E+00 | 1806.96 |
| 412163 | 0.000 | 0.055 | Quinoprotein-ADH-like WSC | *Marssonina brunnea* | EKD19016 | 0.00E+00 | 671.774 |
| 701506 | 0.000 | 0.640 | Tannase and feruloyl esterase | *Oidiodendron maius* | KIM93823 | 0.00E+00 | 677.552 |
| 686719 | 0.350 | 0.966 | Tyrosinase | *Marssonina coronariae* | OWP00047 | 0.00E+00 | 581.637 |
| 579440* | 0.091 | 0.841 | Tyrosinase | *Rhynchosporium commune* | CZT11605 | 8.20E-160 | 461.07 |
| 595981 | 0.000 | 0.492 | Tyrosinase | *Oidiodendron maius* | KIN02411 | 0.00E+00 | 543.117 |
| 50069 | 0.000 | 0.482 | Tyrosinase | *Colletotrichum gloeosporioides* | EQB46731 | 4.10E-148 | 431.409 |
| 657273 | 1.625 | 1.878 | WSCs-galactose binding domain | *Glarea lozoyensis* | XP_008079841 | 1.20E-88 | 301.597 |

<sup>1</sup>Exponentially modified protein abundance indexes (emPAI) (32) after five and seven days growth on ground lodgepole pine. Abbreviations: ADH, Alcohol dehydrogenase; WSC, domain of unknown function possibly involved in carbohydrate binding.
\*Extended protein model 54274 preferred.

Additional extracellular redox enzymes, not classified within the CAzyme system, are listed in Table 3. Perhaps involved in melanin synthesis by DSEs such as *P. scopiformis*, four tyrosinases were identified after seven days incubation. The roles of chloroperoxidases (Phisc_576987; Phisc_588357) and heme peroxidase WSC (Phisc_729673) are more difficult to discern as these enzymes catalyze diverse and often non-specific reactions. The WSC domain (**<u>PF01822</u>**) is widely distributed among fungi and often assumed to be involved in the carbohydrate binding of various glycoside hydrolases and copper radical oxidases. The *P. scopiformis* genome contains at least 38 gene models with WSC domains, of which 15 functionally diverse proteins were identified (five copper radical oxidases, a heme peroxidase, a GH18 chitinase, a GH16 mixed-link glucanase, an AA1 multicopper oxidases and six hypotheticals (Tables 2, 3, and S1 File).

Gene expression data is not yet available for related DSEs, but genome comparisons of Schlegel et al (17) and Knapp et al (18) suggested that the expansion of CAzyme-encoding genes in DSEs genomes, including the root endophyte *P. subalpina*, may confer the ability for saprophytic growth. In this connection, other studies demonstrated that Phialocephala fortinii can hydrolyze carboxymethylcellulose (12, 13) and colonize the tracheids of softwoods (14). In addition, it has been shown that other root-and foliage-associated DSEs will utilize a wide spectrum of substrates (15, 16). Thus, latent saprotrophy may be widespread among DSEs including, at least in the initial stages, the decomposition of forest litter (47), twigs (48, 49) and logs (50). Considering this enormous carbon pool (51), DSEs likely play an important role in the carbon cycle and merit consideration in current ecosystem models (52–54).

## Supporting Information

**S1 File. Detailed information for 590 *P. scopiformis* proteins**. Protein abundance, putative function, CAzyme assignments, InterPro, GO and enzyme codes for 590 P. scopiformis proteins.

## References

1. Eriksson K-EL, Blanchette RA, Ander P. Microbial and enzymatic degradation of wood and wood components. Timell TE, editor. Berlin: Springer-Verlag; 1990.

2. Treseder KK, Lennon JT. Fungal traits that drive ecosystem dynamics on land. Microbiol Mol Biol Rev. 2015;79(2):243–62.

3. Tedersoo L, Bahram M, Polme S, Koljalg U, Yorou NS, Wijesundera R, et al. Fungal biogeography. Global diversity and geography of soil fungi. Science. 2014;346(6213):1256688.

4. Schmidt O. Wood and tree fungi: Biology, damage, protection and use. Berlin: Springer; 2006. 380 p.

5. Riley R, Salamov AA, Brown DW, Nagy LG, Floudas D, Held BW, et al. Extensive sampling of basidiomycete genomes demonstrates inadequacy of the white-rot/brown-rot paradigm for wood decay fungi. Proc Natl Acad Sci U S A. 2014;111(27):9923–8.

6. Hatakka A, Hammel KE. Fungal biodegradation of lignocelluloses. In: Hofrichter M, editor. Industrial Applications. 10. 2 ed. Berlin: Springer-Verlag; 2010.

7. Nilsson T, Daniel G, Kirk TK, Obst J. Chemistry and microscopy of wood decay by some higher ascomycetes. Holzforschung. 1989;43:11–8.

8. Shary S, Ralph SA, Hammel KE. New insights into the ligninolytic capability of a wood decay ascomycete. Appl Environ Microbiol. 2007;73(20):6691–4.

9. Eriksson KE, Blanchette R, Ander P. Microbial and enzymatic degredation of wood and wood components. Berlin: Springer-Verlag. p. 1990.

10. Peay KG, Kennedy PG, Talbot JM. Dimensions of biodiversity in the Earth mycobiome. Nat Rev Microbiol. 2016;14(7):434–47.

11. Mandyam K, Jumpponen A. Seeking the elusive function of the root-colonizing dark septate endophytic fungi. Studies in Mycology. 2005;53:173–89.

12. Caldwell BA, Jumpponen A, Trappe JM. Utilization of major detrital substrates by dark-septate, root endophytes. Mycologia. 2000;92:230–2.

13. Surono, Narisawa K. The dark septate endophytic fungus Phialocephala fortinii is a potential decomposer of soil organic compounds and a promoter of Asparagus officinalis growth. Fungal Ecology. 2017;28:1–10.

14. Sieber TN. Fungal root endophytes. In: Waisel Y, Eshel A, Kafkafi U, editors. Plant Roots: The Hidden Half. Third ed. New york: Marcel Dekker; 2002. p. 887–917.

15. Knapp DG, Kovacs GM. Interspecific metabolic diversity of root-colonizing endophytic fungi revealed by enzyme activity tests. FEMS Microbiol Ecol. 2016;92(12).

16. Carroll G, Petrini O. Patterns of substrate utilixation by some fungal endophytes from conifereous foliage. Mycologia. 1983;75:53–63.

17. Schlegel M, Munsterkotter M, Guldener U, Bruggmann R, Duo A, Hainaut M, et al. Globally distributed root endophyte Phialocephala subalpina links pathogenic and saprophytic lifestyles. BMC Genomics. 2016;17(1):1015.

18. Knapp DG, Nemeth JB, Barry K, Hainaut M, Henrissat B, Johnson J, et al. Comparative genomics provides insights into the lifestyle and reveals functional heterogeneity of dark septate endophytic fungi. Scientific reports. 2018;8(1):6321.

19. Frasz SL, Walker AK, Nsiama TK, Adams GW, Miller JD. Distribution of the foliar fungal endophyte Phialocephala scopiformis and its toxin in the crown of a mature white spruce tree as revealed by chemical and qPCR analyses. Can J For Res. 2014;44:1138–43.

20. Walker AK, Frasz SL, Seifert KA, Miller JD, Mondo SJ, LaButti K, et al. Full Genome of Phialocephala scopiformis DAOMC 229536, a Fungal Endophyte of Spruce Producing the Potent Anti-Insectan Compound Rugulosin. Genome Announc. 2016;4(1).

21. Lombard V, Golaconda Ramulu H, Drula E, Coutinho PM, Henrissat B. The carbohydrate-active enzymes database (CAZy) in 2013. Nucleic Acids Res. 2014;42:D490–D5.

22. Gaskell J, Blanchette RA, Stewart PE, BonDurant SS, Adams M, Sabat G, et al. Transcriptome and Secretome Analyses of the Wood Decay Fungus Wolfiporia cocos Support Alternative Mechanisms of Lignocellulose Conversion. Appl Environ Microbiol. 2016;82(13):3979–87.

23. Hori C, Ishida T, Igarashi K, Samejima M, Suzuki H, Master E, et al. Analysis of the Phlebiopsis gigantea genome, transcriptome and secretome provides insight into its pioneer colonization strategies of wood. PLoS Genet. 2014;10(12):e1004759.

24. Fernandez-Fueyo E, Ruiz-Duenas FJ, Ferreira P, Floudas D, Hibbett DS, Canessa P, et al. Comparative genomics of Ceriporiopsis subvermispora and Phanerochaete chrysosporium provide insight into selective ligninolysis. Proc Natl Acad Sci U S A. 2012;109(14):5458–63.

25. Ryu JS, Shary S, Houtman CJ, Panisko EA, Korripally P, St John FJ, et al. Proteomic and functional analysis of the cellulase system expressed by Postia placenta during brown rot of solid wood. Appl Environ Microbiol. 2011;77(22):7933–41.

26. Nesvizhskii AI, Keller A, Kolker E, Aebersold R. A statistical model for identifying proteins by tandem mass spectrometry. Anal Chem. 2003;75(17):4646–58.

27. Ohtaki S, Maeda H, Takahashi T, Yamagata Y, Hasegawa F, Gomi K, et al. Novel hydrophobic surface binding protein, HsbA, produced by Aspergillus oryzae. Appl Environ Microbiol. 2006;72(4):2407–13.

28. Timell TE. Recent progress in the chemistry of wood hemicelluloses. Wood Sci Technol. 1967;1:45–70.

29. Rytioja J, Hilden K, Yuzon J, Hatakka A, de Vries RP, Makela MR. Plant-polysaccharide-degrading enzymes from Basidiomycetes. Microbiol Mol Biol Rev. 2014;78(4):614–49.

30. Nghi do H, Bittner B, Kellner H, Jehmlich N, Ullrich R, Pecyna MJ, et al. The wood rot ascomycete Xylaria polymorpha produces a novel GH78 glycoside hydrolase that exhibits alpha-L-rhamnosidase and feruloyl esterase activities and releases hydroxycinnamic acids from lignocelluloses. Appl Environ Microbiol. 2012;78(14):4893–901.

31. Levasseur A, Drula E, Lombard V, Coutinho PM, Henrissat B. Expansion of the enzymatic repertoire of the CAZy database to integrate auxiliary redox enzymes. Biotechnol Biofuels. 2013;6(1):41.

32. Ishihama Y, Oda Y, Tabata T, Sato T, Nagasu T, Rappsilber J, et al. Exponentially modified protein abundance index (emPAI) for estimation of absolute protein amount in proteomics by the number of sequenced peptides per protein. Mol Cell Proteomics. 2005;4(9):1265–72.

33. Cowling EB. Comparative biochemistry of the decay of sweetgum sapwood by white-rot and brown-rot fungi. Technical bulletin No, 1258. Washington. D.C.: U.S. Department of Agriculture; 1961.

34. Kleman-Leyer K, Agosin E, Conner AH, Kirk TK. Changes in Molecular Size Distribution of Cellulose during Attack by White Rot and Brown Rot Fungi. Appl Environ Microbiol. 1992;58(4):1266–70.

35. Kerem Z, Hammel KE. Biodegradative mechanism of the brown rot basidiomycete Gloeophyllum trabeum: Evidence for an extracellular hydroquinone-driven fenton reaction. FEBS Lett. 1999;446(1):49–54.

36. Paszczynski A, Crawford R, Funk D, Goodell B. De novo synthesis of 4,5-dimethoxycatechol and 2, 5-dimethoxyhydroquinone by the brown rot fungus Gloeophyllum trabeum. Appl Environ Microbiol. 1999;65(2):674–9.

37. Jensen KA, Jr., Houtman CJ, Ryan ZC, Hammel KE. Pathways for extracellular Fenton chemistry in the brown rot basidiomycete Gloeophyllum trabeum. Appl Environ Microbiol. 2001;67(6):2705–11.

38. Cohen R, Suzuki MR, Hammel KE. Differential stress-induced regulation of two quinone reductases in the brown rot basidiomycete Gloeophyllum trabeum. Appl Environ Microbiol. 2004;70(1):324–31.

39. Daniel G, Volc J, Filonova L, Plihal O, Kubatova E, Halada P. Characteristics of Gloeophyllum trabeum alcohol oxidase, an extracellular source of H2O2 in brown rot decay of wood. Appl Environ Microbiol. 2007;73(19):6241–53.

40. Presley GN, Zhang J, Schilling JS. A genomics-informed study of oxalate and cellulase regulation by brown rot wood-degrading fungi. Fungal Genet Biol. 2016.

41. Westereng B, Ishida T, Vaaje-Kolstad G, Wu M, Eijsink VG, Igarashi K, et al. The putative endoglucanase PcGH61D from Phanerochaete chrysosporium is a metal-dependent oxidative enzyme that cleaves cellulose. PLoS ONE. 2011;6(11):e27807.

42. Yakovlev I, Vaaje-Kolstad G, Hietala AM, Stefanczyk E, Solheim H, Fossdal CG. Substrate-specific transcription of the enigmatic GH61 family of the pathogenic white-rot fungus Heterobasidion irregulare during growth on lignocellulose. Appl Microbiol Biotechnol. 2012;95(4):979–90.

43. Kojima Y, Varnai A, Ishida T, Sunagawa N, Petrovic DM, Igarashi K, et al. A lytic polysaccharide monooxygenase with broad xyloglucan specificity from the brown-rot fungus Gloeophyllum trabeum and its action on cellulose-xyloglucan complexes. Appl Environ Microbiol. 2016;82(22):6557–72.

44. Agger JW, Isaksen T, Varnai A, Vidal-Melgosa S, Willats WG, Ludwig R, et al. Discovery of LPMO activity on hemicelluloses shows the importance of oxidative processes in plant cell wall degradation. Proc Natl Acad Sci U S A. 2014.

45. Bissaro B, Rohr AK, Muller G, Chylenski P, Skaugen M, Forsberg Z, et al. Oxidative cleavage of polysaccharides by monocopper enzymes depends on H2O2. Nat Chem Biol. 2017;13(10):1123–8.

46. Meier KK, Jones SM, Kaper T, Hansson H, Koetsier MJ, Karkehabadi S, et al. Oxygen activation by Cu LPMOs in recalcitrant carbohydrate polysaccharide conversion to monomer sugars. Chem Rev. 2018;118(5):2593–635.

47. Guerreiro MA, Brachmann A, GBegerow D, Persoh D. Transient leaf endophytes are the most active fungi in 1-year-old beech leaf litter. Fungal Divesity. 2018;89:237–51.

48. Boddy L, Griffith GS. Role of endophytes and latent invasion in development of decay communities in sapwood of angiospermous trees. Sydowia. 1989;41:41–73.

49. Griffith GS, Boddy L. Fungal decomposition in ash, beech and oak. New Phytol. 1990;116:407–15.

50. Cline LC, Schilling JS, Meke J, Groenhof E, Kennedy P. Ecological and functional effects of fungal endophytes on wood decomposition. Functional Ecology. 2018;32:484–94.

51. Fraver S, Milo AM, Bradford JB, D’Amato AW, LKenefic L, Palik BJ, et al. Woody debri volume depletion through decay: Implications for biomass and carbon accounting. Ecosystems. 2013;16:1262–72.

52. Wieder WA, Grandy C, Taylor P, Bonan G. Representing life in the Earth system with soil microbial functional traits in the MIMICS model. Geoscientific Model Development Discussions. 2015;8:2011–52.

53. Allison SD, Wallenstein MD, Bradford MA. Soil-carbon response to warmin dependent on microbial physiology. Nature Geoscience. 2010;3:336–40.

54. Shi M, Fisher JB, Brzostek ER, Phillips RP. Carbon cost of plant nitrogen acquisition: global carbon cycle impact from an improved plant nitrogen cycle in the Community Land Model. Global Change Biology. 2016;22:1299–314.

